# Contextual effects and the components of reward sensitivity: Sucrose consumption, discrimination and incentive contrast

**DOI:** 10.1101/2025.08.20.671310

**Authors:** Kristina M. Thompson, Emma L. Stewart, Adam K. Quinn, Howard C Cromwell

## Abstract

Reward sensitivity (RS) has been a moving target because the majority of experimental work utilizes different measures and distinct outcomes in a variety of contexts. Moreover, studies focus on individual components of RS such as intake or reward discrimination or relative valuation. Pinpointing how consistent or variable these distinct components of RS are would be highly beneficial, so we tested identical subjects between two well-established contexts (home cage and operant chamber) using identical sucrose solutions to assess key elements of RS (e.g., intake and reward discrimination and incentive contrast). Previous work has found context invariance for specific components of RS and consistency across contexts could be highly adaptive. Interestingly, the current study found significant discrimination and contrast effects in the home environment and reduced or weaker effects in the operant testing chamber environment. Negative contrast effect was observed only in the homecage and not in the operant chamber. Surprisingly, relative to home, rats consumed significantly more sucrose in the operant box for 1% and 10% but not 30% sucrose. The results support a substantial environmental gating influence whereby context may work to induce alterations in reward value. Contextual factors such as arousal, stress or effort could influence the intensity or expression of RS. Careful consideration to context is highly warranted when generalizing reward processing functions among distinct environmental-experimental settings. Moreover, contextual factors should be recognized as interactive forces that can guide RS as a dynamic state as opposed to a static trait-like indicator of the reward process.

## 1 Introduction

Reward sensitivity (RS) is an individual characteristic and indicators of RS indicate how well the reward process is operating. Measuring RS has been a very useful way to attempt to relate behaviors to future choices and motivation for both natural and drug reward (Stephens et al., 2010). RS has been measured in diverse ways using a wide range of environmental contexts (Toates, 1986; Berridge, 2004; Stephens et al., 2010). Home cage consumption is a staple for intake measures, choice and preference as well as consummatory contrast (Flaherty, 1996). Behavioral chambers typically use operant responses that allow animals access to reward outcomes (Skinner, 1938). In general, assumptions are made that indicators of consumption, discrimination and incentive contrast for specific outcomes would work together among these different environments. For example, a significant preference as discrimination for a higher concentration sucrose solution over a lower concentration should be observed reliably in diverse contexts. This has been explored specifically with consummatory incentive contrast paradigms (Flaherty, Hrabinski and Grigson, 1990). Context invariance has been found in a robust manner in that negative contrast remains following substantive shifts in context and cues used to signal upcoming reward (Grigson, Spector and Norgren, 1993; Daniel et al., 2008). Wiegmann and Smith (2009) manipulated conditioned stimuli and context and found reliable incentive contrast regardless of the shifts in cues or context. This study was done in honey bees (*Apis mellifera*).

Other work has found that incentive contrast is reliable across contexts or using different environmental cues (Flaherty and Avdzej, 1976; Flaherty, Blitzer and Collier, 1987). There has been work showing that incentive contrast can uncouple, specifically when the behavioral responses used to measure the effects differ. Appetitive phase responses sometimes labeled instrumental responses are those related to approach and seeking reward outcomes while consummatory behaviors are the direct responses to outcome acquisition (e.g., licking, intake and ingestion; Craig, 1918). Incentive contrast measured with instrumental action has been shown to be distinct from consummatory measures (Flaherty, 1996). For example, using an alleyway to measure running speed and intake for consumption, incentive contrast was obtained for sucrose intake but not for the instrumental responses (Flaherty and Caprio, 1976). More recent work has found similar dissociations (Sastre et al., 2005). Typically, instrumental contrast is more difficult and expression of it depends more heavily on the learning history, the duration of training, and the outcome used (Webber et al., 2016; Capaldi, 1972; Mellgren, 1971; Panksepp and Trowill, 1971). What is lacking in the experimental work is a study that examines different components of the reward process and RS using the same animals and identical outcomes. Also, a valuable study would include a consummatory condition that attempts to mimic aspects of the instrumental condition with shorter duration outcome exposure using repeated trials instead of a chronic drinking episode (e.g., 30 min to 24-hour exposure).

This study fills an important gap by disentangling specific components of reward processing, namely reward discrimination and relative valuation. These components are important building blocks of preference and can be dissociated and examined outside of standard choice paradigms (Ricker et al., 2016a; Ricker et al., 2016b; Webber et al., 2015). One way to reveal discrimination is by measuring responses during exposure to an outcome in isolation and comparing how these responses differ among alternatives (Crespi, 1942; Ehrenfreund & Badia, 1962; Cleland, Williams, & Dilollo, 1969). This between-session approach to reward discrimination provides a way to evaluate discrimination based on ‘absolute’ reward value of each outcome and removes potential contrast effects. Licking and taste reactions can be used to evaluate discrimination consummatory responses (Panksepp & Trowill,1971; Berridge, 2000; Gutierrez and Simon, 2012) while appetitive measures typically use response rate (Kirkpatrick et al., 2014; Delamater and Nicolas, 2016; Chang et al., 2012) or response latency (Watanabe et al., 2001; Webber et al., 2015; Hermoso-Mendizabal et al., 2020). These effects of discrimination among reward outcomes are reliable during alterations in outcome magnitude, probability, delay or quality. Despite the consistency across diverse studies, reward discrimination can break down due to rearing condition, drug exposure and other variables (Kirkpatrick et al., 2014; Stephens et al., 2010; Ricker et al., 2016a).

Different from discriminating between one outcome and another, reward relativity utilizes a memory of past reward experience and adjusts the value of an outcome depending upon the difference between the previous and present experiences (Flaherty, 1996). In some way, this traditional work could be labeled as reward-gating (Hermoso-Mendizabal et al., 2020) because the behavior is filtered using this reward value adjustment. Incentive contrast (IC) paradigms are often used to measure reward relativity and examine factors involved in the dynamic processes of choice, preference, and decision-making (Webber et al., 2015; Eisenberger et al., 1975). Contrast effects are obtained consistently by measuring consummatory actions such as licking rates and food consumption (Flaherty et al., 1996). Negative contrast (NC) occurs when the value decreases for an outcome relative to previous experience and motivation declines (Weinstein, 1978; Flaherty & Largen, 1975; Flaherty, 1982; Webber et al., 2015). Positive contrast (PC) is the opposite with a relative value increase and an enhancement of motivation (Weinstein, 1978; Flaherty & Largen, 1975; Webber et al., 2015).

Even though incentive contrast effects are quite reliable and observed in diverse species, it can be difficult to obtain in certain contexts and experimental designs. Generally, NC is often found at higher rates than positive contrast (PC; Flaherty, 1982). More successful PC findings have been found when studies implement specific time delayed rewards, deprivation levels, and reward history variables (Flaherty, 1982; Annicchiarico et al., 2016). These effects are sensitive to the dependent measure employed (Papini, 2014). When animals are working for reward outcomes using instrumental or operant behavior (Webber et al., 2015; Zeaman, 1949), contrast effects can be replaced with induction effects (Webber et al., 2015; Weatherly et al., 2006). These induction effects can be expressed in the opposite way compared to contrast with similar rather than different responses among outcomes. Part of this effect is related to generalization between outcomes and seems to rely on the impact of one outcome to dominate and compel the organism to respond in a similar manner to all alternatives. Reward discrimination can be present even when there is an absence of incentive contrast effects (Ricker et al., 2016b). This occurs when responses to distinct outcomes vary; but responses to the same outcome in different reward contexts remain the same.

Reward discrimination and relative valuation should intimately depend upon the individual outcomes and outcome disparity. When these parameters remain constant, components of reward processing should remain stable. Take the outcome and place it in these different contexts and its value moves with it. With this in mind, we utilized identical outcome sets of sucrose concentrations and tested reward discrimination and relative valuation ‘at home’ and ‘away from home’.

## 2. Materials and Methods

### 2.1. Animals

Twelve female rats (Sprague - Dawley strain; >90 days old at start of test sessions) were used. All rats were single housed in 65 × 24 × 15 cm cages; rats ranged in weight from 240 to 320 grams before testing. Female rats were used because they have been neglected overall in behavioral studies and these animals have been found to show higher rates of motivation for sucrose and drug reward. We have related work exploring sex differences in RS using similar paradigm (Thompson et al., 2021). The animals were housed in a colony room on an automatic 12:12 reversed light: dark cycle beginning an hour before testing. The animals were run in four cohorts and given ad libitum (ad-lib) access to food (Harlan Teklad Rat Chow #8604) before testing. Baseline weights were taken by averaging the last three days of recorded weights. Then rats were food restricted to reach a target weight that was 87% of their baseline weights. Once the rats reached their target weights, rats were given about 5 to 15 grams of food each day to maintain this weight. Body condition, hydration status, and maintained food consumption was monitored every day to benefit the heath of the rats. The animals were food restricted for the remainder of this experiment. All procedures were approved by the Institutional Animal Care and Use Committee at Bowling Green State University (protocol 1398631-7).

### 2.2. Sucrose solution

99.5% sucrose powder (Sigma Aldrich, St. Louis MO. Product # S9378) was mixed with tap water (tap water is what all the rats get to drink normally) to create the sucrose concentrations. Sucrose was mixed fresh into 1%, 10% and 30% concentrations every Monday before testing and disposed of every Friday after testing. The solutions were kept in a mini fridge only available to the lab (not used for storage of any other food, drinks, or substances) that is located between the housing room and testing room. The making of sucrose and storage of the sucrose in the mini fridge ensured the quality of the sucrose concentrations and prevented sickness from any build up or breakdown of the sucrose molecules in the tap water.

### 2.3. Training and Testing Procedure in the Home-cage

During training, rats were transported in their home cages (rats were not removed from their home cage during training or testing for home cage conditions) to a separate red-light testing room and presented with 30 second trials, with 1-minute intervals, of only 10% sucrose. The separate room was decidedly used to limit rats from viewing each other receiving the sucrose solutions. This would also mimic their solo experiences when moved to the testing room with the operant boxes. The rats had 10 trials of 30 seconds which resulted with a total of 300 seconds of access per session. The sucrose concentrations were administered by manually lowering the syringes (containing 5 ml of sucrose concentration) into the upper left-hand side of their home cages.

During testing, counterbalancing of the daily schedule was done to eliminate predictability of sucrose concentrations. Along with counterbalancing of the daily schedule, the weekly schedule was broken down into two categories: discrimination days and contrast days (see Figure 1A). For discrimination days rats received either 1%, 10% or 30% sucrose concentrations (example trials: 1, 1, 1, 1… or 10, 10, 10, 10….). On contrast days rats were tested specifically for IC by presenting alternating trials of either 1% and 10% or 10% and 30% sucrose concentrations (example trials: 10, 1, 10, 1…) with successive presentation of the syringes contained sucrose solution.

**Figure 1.**
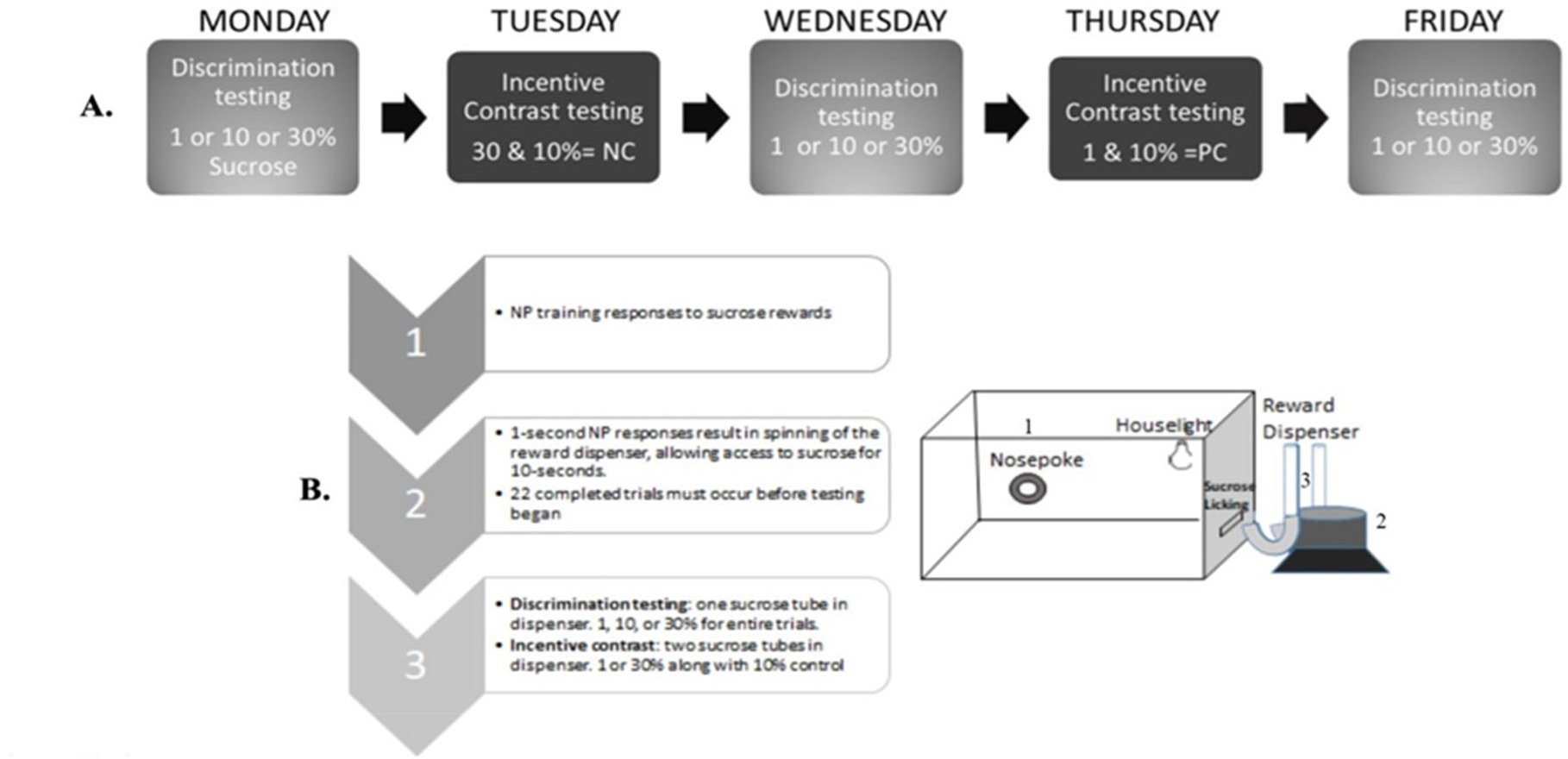
Weekly schedule and apparatus: (A) An example of the counterbalanced daily schedule for a week of testing. The schedule is broken into two categories: discrimination days and contrast days. For discrimination days rats received either 1,10 or 30% sucrose concentrations. On contrast days concentrations were alternated from either 1 or 30% to 10% (control). (B) An example of the operant box apparatus used for training and testing with labels and brief descriptions.

### 2.4 Behavioral training and testing procedures in operant box

After home cage training, the same animals used in behavioral training and testing within standard operant boxes (35 × 30.5 × 30 cm; Med Associates Inc., Fairfax, VT, US) encased in sound-attenuating chambers and connected via software (MED-PC IV) that was used to run the behavioral sessions, deliver reward, and collect data (see Figures 1 A and 1B). The daily sessions each week alternated discrimination testing (e.g. a single outcome) with incentive contrast testing (e.g., alternating outcomes). Each operant box contained a nose poke aperture, lever, house light, camera to observe rat behavior, and a hole that allowed enough space for animals to lick a sucrose solution from a ball-bearing nozzle connected to a graded syringe tube (ml). Tubes containing sucrose solution were attached to an automated turntable outside the boxes that spun after each response and stopped to align the nozzle with the open slot to allow animals to consume the solution. A house light turned on to cue the presence of the solution and the beginning of a trial. At the end of a trial, the turntable spun back to prevent access to the solution and the house light turned off to cue the end of a trial.

Animals were shaped to lick for the sucrose in the operant chamber with access to sucrose available only after a nose poke that led to the house light turning on during a 10 second access period. Identical tubes and spouts that were used in the homecage were used in the operant chamber. For the first three days of training the tubes were hand-held and placed through a slot in the right-hand side of the operant box, paired with a light, so that the rats associated their nose pokes with the tube of sucrose. After the three days, the tubes were connected to the rotating plate/tube device designed and manufactured in the Department of Psychology, BGSU (see Figure 1B).

If a successful nose poke occurred a light attached to the top of the operant chamber would turn on and the rotating plate would spin the tube to the small opening in the right-hand side of the operant box. The opening was small enough to only allow the rats to lick through it or stick their nose through but not at the same time. In order to receive their reward, the rats had to stick their tongues through the slit to reach and proceed to lick the tube. During training the rats had 30 seconds, from the time the light turns on, to lick as much as they wanted of 10% sucrose. After the 30 seconds the light would turn off and the spinning plate would spin the tube away from the slit so that the tube was inaccessible to the rat. Once the rats were consistently completing 22 trials, testing sessions commenced.

During testing the rats only had 10 seconds, from the time the light turned on, to lick as much as they wanted. After the 10 seconds the light turned off and the spinning plate spun the tube away. For testing, rats had to complete 22 trials in 30 minutes; once 22 trials were done the rats were removed from the operant box (even if the full 30 minutes had not yet passed). Overall the rats that completed all 22 trials at 10 seconds would have had a total of 220 seconds of sucrose access per session. The rats were presented with the sucrose with the same counterbalanced daily schedule as in the home cage. On discrimination days rats were only given one type of sucrose, thus the spinning plate was only equipped with one tube containing that specific solution. On contrast day rats were given two sucrose concentrations, thus the spinning plate was equipped with two tubes which spun so that the tubes alternated each nose poke.

### 2.4. Statistical Analysis

Master data spreadsheets (Excel, Microsoft Inc.) were used to organize and report daily weights, concentration(s) of sucrose, the time their trial began and ended, and all trial consumptions. Sucrose consumed was converted to grams/kilogram for each concentration. Two weeks of testing were recorded however only the second week of data was used for analysis reported due to increase in familiarity with the solutions and greater reward sensitivity shown by the rats during these weekly sessions. Statistical Package for the Social Sciences (IBM SPSS Ver. 24) was used for all statistical data analysis. For the home cage consummatory contrast, the amount of sucrose consumed was documented for each trial. Non-parametric analysis on the amount consumed as well as the average consumption for each subject was calculated. An Friedman’s ANOVA with contrast condition (PC vs. Control or NC vs. Control) as main factor followed by pairwise tests were completed. Similar analysis was completed for discrimination among the three solutions using an Friedman’s ANOVA with three levels for the sucrose solutions (1, 10 and 30%) followed by pairwise tests.

During testing in the operant chamber, consumption was recorded manually after session end in milliliters (mL) and converted to g/kg/trial. For discrimination, we examined differences in consumption volume among the three concentrations. For incentive contrast, we examined differences in consumption of the 10% sucrose depending upon the alternative outcome (1 or 30%). Behavioral data for nose-poke latencies and licking rates were acquired using MED-PC software (Med. Associates Inc., VT, USA). Each lick was recorded by a custom written program. Lick rates (LR) were obtained as an average from the 10s access time to the sucrose solution. Nose poke (NP) latency measures were the time between one nose poke and the next nose poke that initiated a trial. This latency necessarily includes the 10s sucrose access period as well. Custom-written programs recorded latencies and nose-pokes to allow for complete analyses. Latencies and lick rates from the trial set were averaged for each animal and grouped according to session type. Non-parametric analysis was performed on each dependent measure (average nose-poke latency and licking rates) using the 2 × 2 ANOVA with sex and contrast-control conditions as the main factors. Data was explored using descriptive statistics, Q-Q plots, and tests for normality (e.g., Shapiro-Wilks test) for use of non-parametric statistical analysis. Nose pokes made during the lick access period were considered errors because of being non-contingent to the sucrose acquisition. These nose poke errors were analyzed due to seeing multiple occurrences of them during testing sessions in the operant box.

## 3. RESULTS

### 3.1. Discrimination in the Homecage

To determine how reward processing differs between homecage and operant chamber contexts, it must first be determined how behavior occurred within each context. The distribution of 1%, 10%, and 30% sucrose consumptions (g/kg) within the homecage were compared to determine if the rats could discriminate between each concentration (Figure 2A). Friedman’s two-way analysis of variance indicated that the distributions were significantly different from each other, *F (2) = 22, p =*.*001*. Wilcoxon tests were then used to determine the median difference between each concentration. Analysis indicated that the rats significantly consumed lower amounts of 1% (*M =*.*01, SD =*.*01*) compared to 10% (*M=*.*53, SD =*.*05*), *z = 3*.*06, p =*.*002*. They significantly consumed a lower amount of 10% compared to 30% (*M = 1*.*79, SD =*.*21*), *z = 2*.*94, p =*.*003*. Lastly, they consumed significantly more 30% compared to 1%, *z = 3*.*05, p =*.*002*.

**Figure 2.**
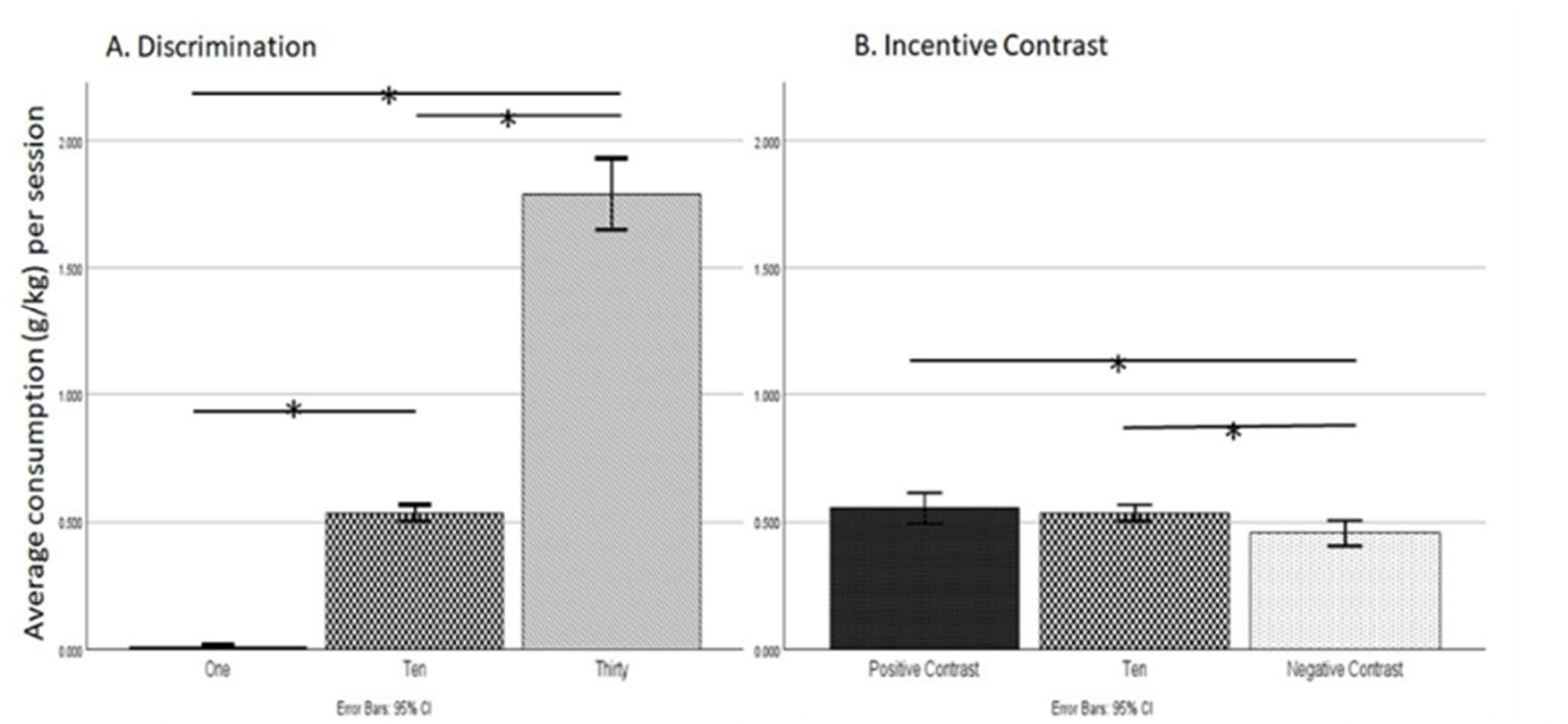
Discrimination and Incentive Contrast average consumptions rates within the Homecage: (A) Discrimination: rats significantly consumed higher amount of 30% compared to 10% and 1%. They also significantly consumed higher amounts of 10% compared to 1%. * p <.05. (B) Incentive contrast: rats significantly consumed a higher amount when they were upshifted (PC) compared to downshifted (NC), they also consumed more baseline 10% compared to the downshift (NC).* p <.05.

### 3.2. Incentive Contrast (IC) in the Home Cage

For positive and negative contrast, the baseline 10% consumption rate was compared to the 10% consumption rate **after** upshifts from 1% (PC) or downshifts from 30% (NC) (Figure 2B). Friedman’s two-way analyses indicated that the distributions between the baseline 10% and the shifts (PC and NC) were significantly different, *F (2) = 11*.*46, p =*.*003*. Wilcoxon tests determined that rats significantly consumed a higher amount of the baseline 10% (*M =*.*53, SD =*.*05*) compared to when there was a down shift, NC 10% (*M =*.*46, SD =*.*07*), *z = −2*.*22, p =*.*026*. The test also determined that the rats significantly consumed more when upshifted, PC 10% (*M =*.*55, SD =*.*09*) compared to when they were down shifted, NC 10%, *z = −2*.*93, p =*.*003*. However, rats did not significantly differ in their consumptions between baseline 10% and the upshifted PC 10%, *z =*.*267, p =*.*79*.

### 3.3. Discrimination in operant chamber

Within the operant chamber, Friedman’s two-way analyses among the distributions of 1%, 10%, and 30% sucrose consumptions (g/kg) was conducted to determine concentration discrimination (Figure 3A). The analysis indicated that the rats could significantly discriminate between the concentrations, *F (2) = 22, p =*.*001*. Wilcoxon tests confirmed that rats significantly consumed a lower amount of 1% (*M= 3*.*26, SD = 1*.*29*) compared to 10% (*M = 4*.*19, SD =*.*84*), *z = 3*.*06, p =*.*002*. Rats also significantly consumed a lower amount of 1% compared to 30% (*M = 4*.*00, SD =*.*75*), *z = 2*.*94, p =*.*003*. Differing from the homecage, rats consumed a significantly higher amount of 10% compared to 30%, *z = 2*.*04, p =*.*03*.

**Figure 3.**
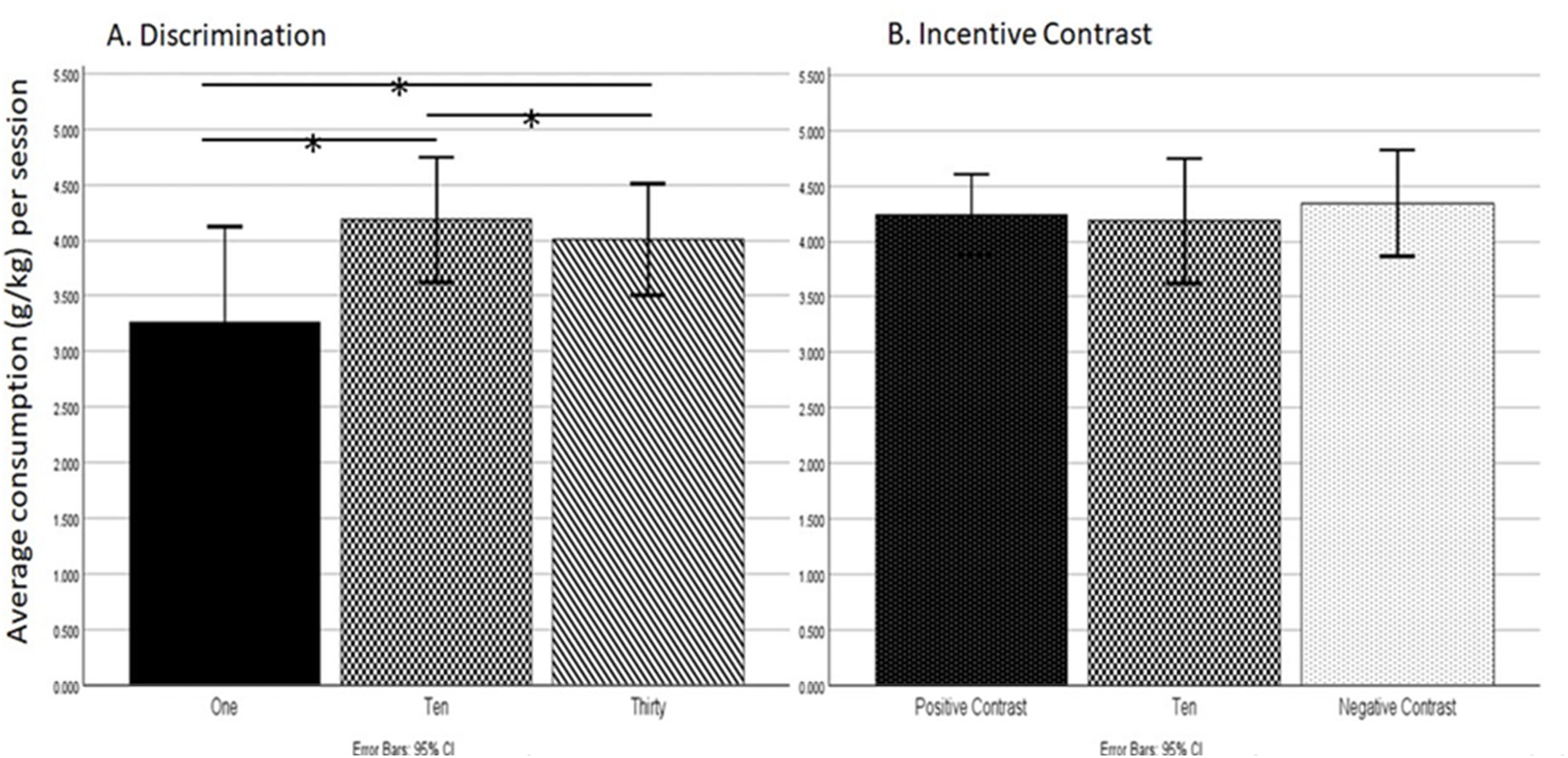
Discrimination and IC Average Consumption Rates within the Operant Box. (A) <u>Discrimination</u>: rats significantly consumed a higher amount 10% compared to both 1% and 30%. Rats also significantly consumed a higher amount of 30% compared to l%.*p <.05. (B) <u>Incentive Contrast</u>: rats did not significantly show changes between baseline 10% and 10% after upshift from 1% (PC) or downshift from 30% (NC).

### 3.4. Incentive Contrast in the Operant Chamber

Similarly, to the homecage, positive and negative contrast were compared by using the baseline 10% consumption rate compared to the 10% consumption rate **after** upshifts from 1% (PC) or downshifts from 30% (NC) (Figure 3B). Friedman’s two-way analyses indicated that the distributions between the baseline 10% and the shifts (PC and NC) were not significantly different, *F (2) = 1*.*27, p =*.*53*.

### 3.5. Licking Rate Discrimination in the Operant Box

Another way to determine if the rats were discriminating between the concentrations within the operant box is to analyze the number of licks they made per trial. The distribution of licking rates for the 1, 10, and 30% sucrose concentrations were compared to determine if the rats were discriminating in their number of licks between each concentration (Figure 4). Friedman’s analysis indicated that there was a significant difference in licking rates between the sucrose concentrations, *F (2) = 6*.*55, p =*.*038*. However, Wilcoxon analysis determined that there was only a significant difference between 1% and 10% licking rates. The rats licked significantly less times for 1% (*M = 32*.*60, SD = 12*.*90*) than 10% (*M = 41*.*86, SD = 8*.*36*), *z = 58, p =*.*026*. There was no significant difference between 10% and 30% licking rates or between 1% and 30%.

**Figure 4.**
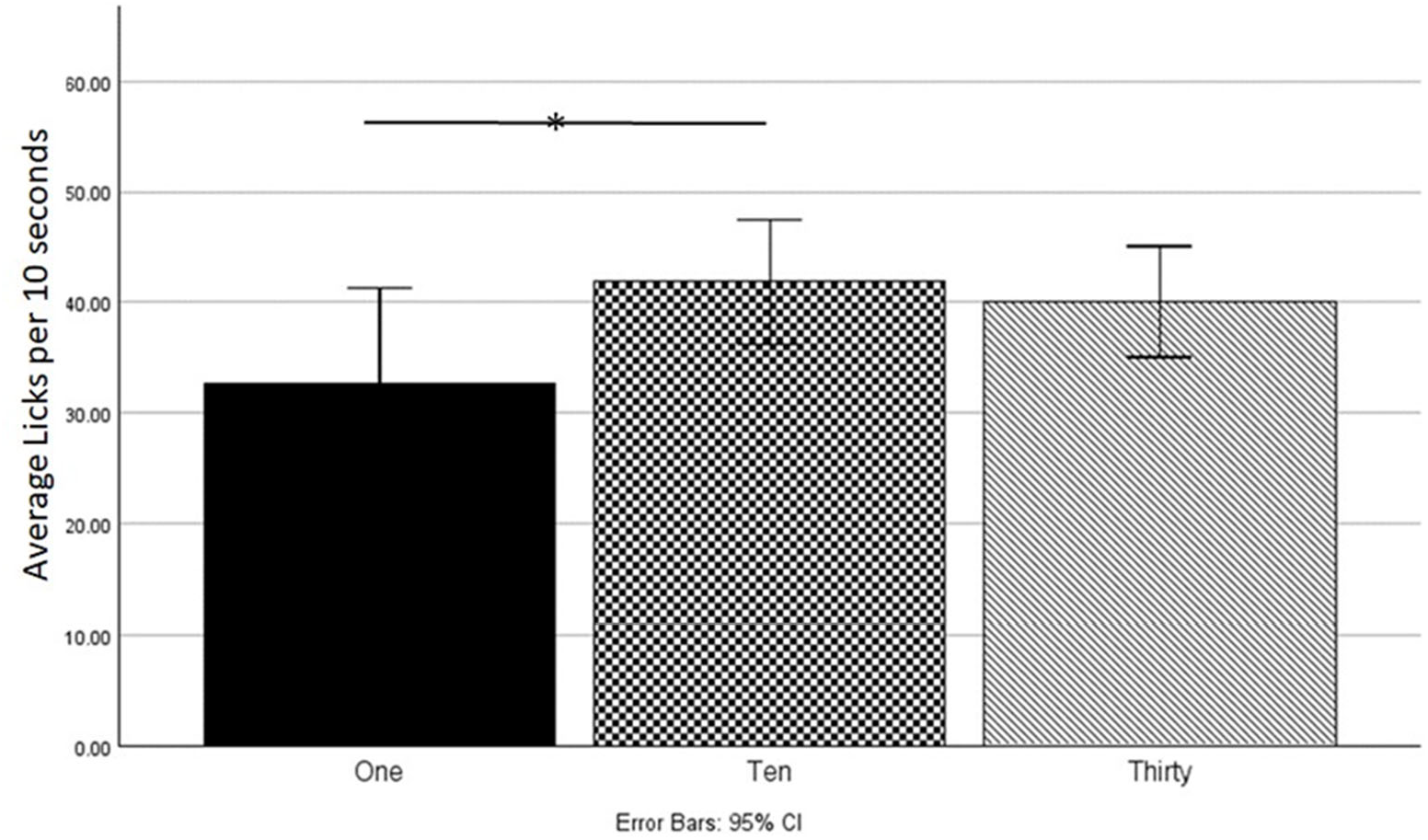
Licking Rate Discrimination within the Operant Box: Discrimination within the average licking rates per 10 seconds of access between 1,10 and 30% concentrations. Rats significantly performed a higher amount of licks per 10 seconds for 10% than 1% but not for any other conditions. *p <.05.

### 3.6. Licking Rate Incentive Contrast in the Operant Box

Analysis was also conducted to determine if the rats could show incentive contrast shifts within their licking rates. Three tests were conducted to determine this; firstly, it was tested to see if their licking rates changed in the first trial after a shift. Freidman’s analysis indicated that there was no significant difference in the rats licking rates between the previous trial licks and the <u>first</u> trial after the shift (positive or negative) was made, *F (2) = 4*.*23, p =*.*120*. The second test was to analyze if there was a significant difference within <u>9</u> trials after a shift; Friedman’s analysis indicated no significant difference in licking rates within 9 trails after a shift, *F (2) = 1*.*27, p =*.*529*. Lastly, a Freidman’s analysis was run to determine if there was significant difference in the number of licks in <u>all</u> trails after a shift. Results indicated there was no significance difference in the total number of licks after a shift, *F (2) =*.*545, p =*.*761*.

### 3.7. Nose Poke Latencies Operant Box

Lastly, the total number of nose pokes and the distribution of nose poke latencies for the 1, 10, and 30% concentrations were compared to determine if the amount of nose pokes and/or the latency between nose pokes indicated discrimination. Friedman’s analysis indicated that the rats did not significantly differ in the total number of nose pokes per trial, *F (2) = 5*.*64, p =*.*06*. Similarly, rats did not significantly differ in latencies nor significantly produce nose pokes after a positive or negative contrast shift, *F (2) = 4*.*23, p =*.*120*. The average nose poke latency for each concentration is as follows: 1% (M = 30.22, SD = 19.37), 10% (M = 18.11, SD = 8.09), 30% (M = 22.62, SD = 10.58), NC (M = 21.80, SD = 11.55), and PC (M = 19.56, SD = 9.15).

## 4. Discussion

Reward discrimination and relative valuation are consistent effects in behavioral sciences and experimental psychology (Young, 1961; Bolles, 1967; Flaherty, 1996); however, there are clear ways to dissociate or impair these abilities. Pre-exposure to drugs of abuse alter reward sensitivity to natural rewards such as food - including sucrose (Stephens et al., 2010). Reward sensitivity can be inherently altered as seen in animals selectively bred for high alcohol intake (McGraw et al., 2024). Other work has found that subjects can have difficulty in reward discrimination and other measures of reward sensitivity due to the parameters of the operant procedure; this includes: the rate of reinforcement, effort, and probability of the outcome (Ricker et al., 2016b; Mellgren, 1972; Rosen, 1966). Certainly all these parameters shift reward value, from the contextual-side, and most work has examined the breakdown of reward sensitivity after altering outcome value directly via changes in reward type, magnitude or access (Spear and Hill 1965; Flaherty and Largen, 1975; Ricker et al., 2016a; Ricker et al., 2016b); however, the nature of relative reward processing is complex and IC paradigms have been inconsistent.

The current study found clear differences for both reward discrimination and relative valuation between contexts with changes in reward type. These changes were obtained despite the use of the exact same outcomes and outcome disparities in both environments. In the home environment, during food restriction, both reward processing components were intact and fit the predictions according to the law of magnitude and incentive contrast effect (Flaherty, 1996; Young, 1961). The effect in the homecage was only obtained in restricted animals so we tested the same condition in the operant chamber. When placed in the operant chamber, reward discrimination and relative valuation broke down.

### 4.1 Contextual influences

By interpreting the environmental-gating of the present work, we can focus on several differences between the environments including some aspects of the task that necessarily differ due to shifts in context. Greater general arousal or stimulation in the operant session could alter key aspects of learning and attention involved in motivated action (Starling et al., 2013; Robbins, 1997). The greater arousal could play a role in the breakdown in reward processing leading to generalized responses to outcomes discriminable in another context (Riemer et al., 2016). Moreover, training effects can lead to reward insensitivity (Dickinson, 1985), leaving animals displaying similar responses among contexts including similar responses between extinction and non-extinction contexts (Perez and Dickinson, 2020).

Our training was relatively minimal for this nose poke-sucrose licking task compared to other operant paradigms, yet even relatively minimal training has been found to embed habit formation (Halverstadt and Cromwell, 2018). Relatedly, reward sensitivity has been linked with changes in primary affective states in animal models (Papini et al., 2006; Glueck et al., 2018) but the relation can be tenuous (Binkley et al., 2014). Animals tend to express greater negative contrast effects following a negative experience such as isolation (Burman et al., 2008) but this effect depends on the way incentive contrast is measured (Mitchell et al., 2012; Binkley et al., 2014). Affective states certainly vary in different contexts with greater anxiety in the operant chamber, a relatively less familiar environment compared to homecage and one with unpredictable outcomes relying on self-paced free operant responding.

The anticipation of each context could also alter the results of contrast; King and colleagues (2002), as well as Webber and colleagues (2015), have indicated that the anticipation for higher value outcomes can cause induction. Positive induction relies on the anticipation for a future higher value outcome (and vice versa for a negative induction), which can possibly alter the dynamic of the alternating outcomes resulting in outcome generalization. While we counterbalanced for each given day, the number of trials and time access did not change and thus (over the course of each session) rats could have anticipated the next outcome within the trial-by-trial session within the operant box. This could have possibly resulted in a breakdown in reward processing leading to generalized responses and the lack of discrimination and IC results that we found.

Interactive with the differences in arousal, the operant chamber also necessarily includes greater effort in obtaining the reward outcome and these changes in effort can alter the reward value (Clement et al., 2000; Knauss et al., 2020). In the homecage, the animals merely had to approach the lick tube and licking behavior was targeting a spout lowered down into the cage. In contrast, the operant chamber contained a slot the animals had to extend the tongue and lick the sucrose solution from a tube outside the cage. Moreover, the animals had to learn to nose poke in a distinct location in order to gain access to the reward. These factors require greater motivation, associative learning, flexible action strategies, and different motor requirements to fit into the distinct categories of consummatory and appetitive behavior (Tinbergen, 1951; Craig, 1917). The bivariate regressions within our analyses between home cage and operant testing demonstrated that there were no significant correlations between their home cage consumption rates and their operant consummatory or behavioral results. Thus, it could be confirmed that the motivational shift in the operant box is more demanding and inadequate, in this instance, to display appropriate discrimination and IC. Previous research has also demonstrated this inadequacy; despite the power of effort to sway choice, preference and reward value, Weatherly and colleagues (2006), found that it was not sufficient to alter induction in an operant task.

The present study differed from previous works which demonstrated reward sensitivity (Crespi, 1942; Ehrenfreund & Badia, 1962; Pieper & Marx, 1963; Cleland et al., 1969; Flaherty & Caprio, 1976; Fisk & Cohen, 1977; Flaherty 1996) by observing both consummatory and instrumental behaviours with a within-session contrast experience and a between session experience. Even though we didn’t see reward sensitivity between contexts, similar work between consummatory and instrumental behaviors by Webber and colleagues (2015) demonstrated and obtained positive and negative IC in sessions with the greatest outcome differential and in initial NP latencies that followed auditory cues to the same outcomes. Overall, we found that reward discrimination and contrast occurred as differences in consumption in one context but not the other. This finding provides a novel separation between consummatory ‘at home’ and consummatory ‘away from home’ measures. This suggests that these actions rely heavily on preceding appetitive behaviors and can be impacted by these previous behaviors in terms of reward outcome processing as also demonstrated by Pieper and Marx (1963). It is possible that the functional nature of reward sensitivity may be dependent upon reward discrimination (as seen in our home cage sensitivity) but independent to reward value (demonstrated by diminished operant context).

### 4.4. Limitations

The study had limitations including the alteration of the order of sucrose presentations between the contexts. We wanted to utilize discrete trial presentations in both contexts but used interspersed trials in the home-cage while using a single shift trial procedure in the operant chamber. Order of reward presentation can have a substantial impact the way outcomes are compared (Douglas et al., 2018). Also, the rats within the homecage experiments did not have the same extensive period of exposure to sucrose as compared to the following operant chamber experiments – this is due to the same rats being used over time in both experiments. This could potentially have led to a more generalized effect within the operant box as the rats grew more accustomed to consuming the different sucrose concentrations. Home cage housing conditions were also not explored in this experiment but could have had an impact on the rat’s behavior (Denny, 1975; Burn, Peters, Day, & Mason, 2006; Wood et. al., 2006; Mitchell et al., 2012). In 2006, Wood and colleagues observed that rats who had previous exposure to high concentration of sucrose drank significantly less when switched to a low concentration in combination with poor housing conditions (wire-bottom cages). Mitchell and colleagues (2012) also demonstrated that home cage environment can have an effect on incentive contrast with animals housed in a barren environment showing faster baseline response times and negative contrast.

There was also a limitation in the use of sexes. Previous work has obtained more consistent contrast and discrimination in the operant setting by using interspersed trials with Sprague-Dawley male rats (McGraw, 2018). Female animals were used in the present work, and this limits the generalizability of the findings. Male rats have been found to emit incentive contrast more consistently compared to female rats (Weinstein, 1978) and female rats have been found to show greater effects in response to novelty and stress (Bangasser and Wicks, 2017). This could lead to male rats responding more similarly among the contexts, yet we have preliminary evidence that males respond overall in a similar fashion with comparably better discrimination and contrast in the homecage using an identical paradigm (Thompson et al., 2021).

### 4.5. Conclusions

In general, it may be that reward sensitivity is not only impaired when moving from one context to another but instead organisms are shifting strategies dependent upon different integrative factors (Fisk & Cohen, 1977; Palminteri et al., 2015). Findings appear as insensitivity based on standard measures, but in actuality the output is optimizing motivation based on interactions among external and internal variables. As Killeen stated in 1993, ‘scales of value determined in one context are likely to suffer nonlinear distortions when employed in another’ (Killeen et al., 1993, p. 213). This makes preferences difficult to track from one context to another because it appears that when rewards are moved between contexts, the value can change significantly. The value learned and used can translate poorly between contexts (Holmes and Westbrook, 2017), and an appreciation of the intrinsic and extrinsic influences of context, and how these interact will lead to better understanding of basic decision-making and motivation based on shifts in valuation of outcomes.

Applications for examining drug effects and addiction are abundant. Future work could explore environmental gating (see Badiani, Caprioli & De Pirro, 2019) effects using identical drug outcomes/doses between ‘home’ and ‘away from home’ outside of using standard concurrent choice paradigms such as between-session discrimination and incentive contrast. One fruitful area includes ethanol intake and preference. Ethanol intake is typically studied in both the homecage and operant environment using similar range of doses (Rodd-Henricks et al., 2002; Samson and Czachowski, 2003) but consumption and motivation can vary significantly between these contexts. Moreover, the results can guide future work exploring neural and psychological mechanisms involved in gating contextual information based on influences from ‘home’ versus ‘away from home’.

